# Chromatin states coordinate diurnal gene expression in the *Drosophila* retina

**DOI:** 10.64898/2026.10.16.759625

**Authors:** Gaoya Meng, Sarah E. McGovern, Lucas D. Kramer, Ricardo Iván Martínez Zamudio, Vikki M. Weake

**Affiliations:** Department of Biochemistry, Purdue University, West Lafayette, Indiana 47907, USA; Purdue Center for Cancer Research, Purdue University, West Lafayette, Indiana 47907, USA; Cold Spring Harbor Laboratory, Cold Spring Harbor, NY, USA; Department of Pharmacology, Rutgers-Robert Wood Johnson Medical School, 675 Hoes Lane West, Piscataway, NJ, USA

**Keywords:** Diurnal rhythms, circadian clocks, epigenomics, transcription, Drosophila, retina, histone modification

## Abstract

Daily biological rhythms enable organisms to anticipate and respond to predictable diurnal environmental changes. While the circadian clock maintains rhythms under constant conditions, it remains unclear what gates the time-of-day expression of genes that are rhythmic under light:dark cycles. In the retina, restricting biological processes to specific times of the day is crucial for homeostasis because disrupting these rhythms causes retinal degeneration. We show that in light-sensitive photoreceptor neurons of *Drosophila*, the circadian transcription factors only account for the timing of a minority of rhythmic genes. Instead, integrative analysis of the daily chromatin landscape and transcription factor footprints reveals that promoter, enhancer, or repressive signatures strongly correspond with phase. Further, genetic modulation of the repressive and promoter chromatin states altered gene expression in a phase-specific manner. Together, our data support a model where the genome is organized into discrete chromatin signatures that collaborate with transcription factors to restrict diurnal transcription to distinct phases of the day.

## Introduction

Daily biological rhythms are maintained by a molecular circadian clock that is largely conserved from *Drosophila* to humans ^1^. Environmental cues or *zeitgebers* such as light entrain this molecular clock by resetting the transcription-translation feedback loop ^1,2^. At the heart of the molecular clock are two basic helix-loop-helix (bHLH) transcription factors, Clock (Clk) and Cycle (Cyc), which heterodimerize and bind to E-box motifs upstream of clock output genes ^1^. Clk:Cyc-activated genes include those encoding their own repressors, *period* (*per*) and *timeless* (*tim*), generating a negative feedback loop with a period of ∼24 h ^1^. In flies, clock neurons in the brain act as the central pacemaker, analogous to the suprachiasmatic nucleus (SCN) in mammals ^1^. This central clock provides synchronizing cues to peripheral clock cells, including the photoreceptor neurons in the retina, which in turn contribute to light entrainment of the central clock ^3^. Although the molecular clock is required to maintain rhythms in the absence of zeitgebers, environmental cues also strongly contribute to daily rhythms ^1^. In the retina, light is a major zeitgeber; 10-fold more genes in the mouse retina exhibit rhythmic expression under light:dark versus constant darkness ^4^. Dietary intake plays a similarly important role: dietary restriction doubles the amount of circadian transcripts in whole flies and delays visual senescence in a Clk-dependent manner ^5^. Thus, environmental cues interact with the molecular clock to dictate daily biological rhythms; here, we define those rhythms that occur under standard light:dark cycles as “*diurnal rhythms*”, in contrast to “*circadian rhythms*” that persist under constant darkness.

The retina is an integrated circadian system that receives multiple environmental inputs to coordinate robust oscillations in its morphology, physiology, and function ^6^. Many retinal biological processes show diurnal rhythms, occurring at specific times of the day with respect to the light cycle, including the retinal light response and sensitivity, synaptic structure, and susceptibility to light damage ^6^. Diurnal rhythms are critical for retinal homeostasis because genetically disrupting the molecular clock affects retinal development and accelerates retinal degeneration. In mice retinal progenitor cells, knockout of *Bmal1* (ortholog of *cyc*) impairs retinal neurogenesis ^7^ and vascular development ^8^. In *Drosophila*, photoreceptor-specific expression of dominant negative Clk results in light-dependent retinal degeneration ^5,9^ with similar degeneration observed in *per^01^* flies ^10^. In mice, retina-specific knockout of *Bmal1* accelerates the age-dependent loss of cone viability and function ^11^. Even mild defects in Per2 (ortholog of *per*) in homozygous *Per2^luc^* mice lead to premature aging of photoreceptor cells and retinal pigment epithelium ^12^. Clock disruption also exacerbates inherited retinal dystrophies because the rod-specific *Bmal1* knockout increases degeneration in mouse models of retinitis pigmentosa ^13^. Notably, diurnal rhythms can also be extensively rewired without loss of the core clock itself. During aging ^14–17^, neurodegenerative disease ^18^, and altered nutritional states ^19,20^, many genes lose, gain, or shift rhythmic expression, while core clock genes remain largely intact and oscillate normally, a phenomenon termed *circadian reprogramming* ^19,20^. Such reprogramming is widely present in the retina of aged flies and diabetic mice ^10,21–23^. Therefore, maintaining healthy diurnal rhythms in the retina is critical, and requires additional mechanisms to the molecular clock.

Although the molecular clock components have been extensively characterized over the past few decades ^1^, an outstanding question remains: how is diurnal gene expression organized into different time windows across the 24-hour day? The circadian transcription factors cannot account for the varied phases (timing of peak gene expression) of the diurnal transcriptome. This is evident because BMAL1 in mice and Clk:Cyc in flies, show synchronized DNA binding at specific times of the day (ZT4 in mice, ZT12 in flies) across all of their binding sites, but the timing of this transcription factor binding is largely uncoupled from the phase of most target gene expression ^24–27^. Although cooperation between BMAL1 or Clk:Cyc and tissue-specific transcription factors contributes to determining which genes are rhythmic in a given tissue ^26,28^, the impact of this collaboration on the phase of expression is limited. Even the phase of cyclic enhancer RNA, regulated by circadian transcription factors, correlates with only a portion of pre-mRNA phases ^29,30^, suggesting additional mechanisms regulate the timing of expression. Here, we explored the mechanisms that regulate the phase of rhythmic gene expression across the genome in a single, light-responsive cell type – photoreceptors in *Drosophila*. We show that while Clk and Cyc contribute to rhythmicity of about half of diurnal genes in photoreceptors, their occupancy is uncoupled from the phase of transcription for most of their bound genes – the molecular clock genes are a notable exception. Instead, we find that most diurnally rhythmic genes exhibit distinct repressed, promoter, and enhancer chromatin signatures that correspond and contribute to their expression phase. Using ATAC-seq footprinting, we identified transcription factors that are preferentially enriched at promoters or enhancers of genes expressed at specific phases. In contrast, the repressive state lacked these phase-enriched transcription factors, and may be less dependent on transcription factors for transiently low-level expression. Together, our data support a model in which the genome is organized into discrete chromatin signatures that collaborate with transcription factors to restrict diurnal transcription to distinct phases of the day. This model provides a framework to understand and predict how disruptions to the epigenome would result in circadian reprogramming during aging and pathological conditions.

## Results

### Rhythmic transcription is organized into waves across the diurnal cycle

To explore the mechanisms underlying the timing of transcription in *Drosophila* photoreceptors, we enriched GFP-tagged photoreceptor nuclei and profiled the diurnal transcriptome and epigenome of young (10 days post-eclosion) flies. We generated H3K27ac, H4K16ac, and H3K27me3 CUT&RUN profiles at six time points across a 12-h light:12-h dark cycle, and integrated them with our published nuclear RNA-seq, ATAC-seq, and H3K4me1/2/3 data from the same time points ^10^ (**Figure 1A-B**). Time points are indicated in zeitgeber time (ZT) with ZT0 marking lights on. Because our photoreceptor transcriptome data derive from nuclear RNA-seq, rhythmicity parameters from intronic reads and exonic reads were highly correlated (*r*^2^ *= 0.9*) ^10^, indicating these data reflect pre-mRNA rather than steady-state mRNA levels. In young photoreceptors, about half of all expressed genes exhibited rhythmic transcription by dryR analysis (**Figure 1C, Table S2**). These rhythmic genes fell into three broad phase waves in the morning (ZT0-8, n=2558), afternoon (ZT8-16, n=1471), and late night (ZT16-24, n=1588, **Figure 1D**). This triphasic pattern persisted under more stringent expression and rhythmicity cutoffs, indicating that these transcription waves do not arise from lowly expressed genes or rhythm-detection noise (**Figure S1**). A similar triphasic pattern was observed in the baboon retina, although many other tissues exhibited predominantly unimodal or bimodal phase distributions ^31^.

**Figure 1:**
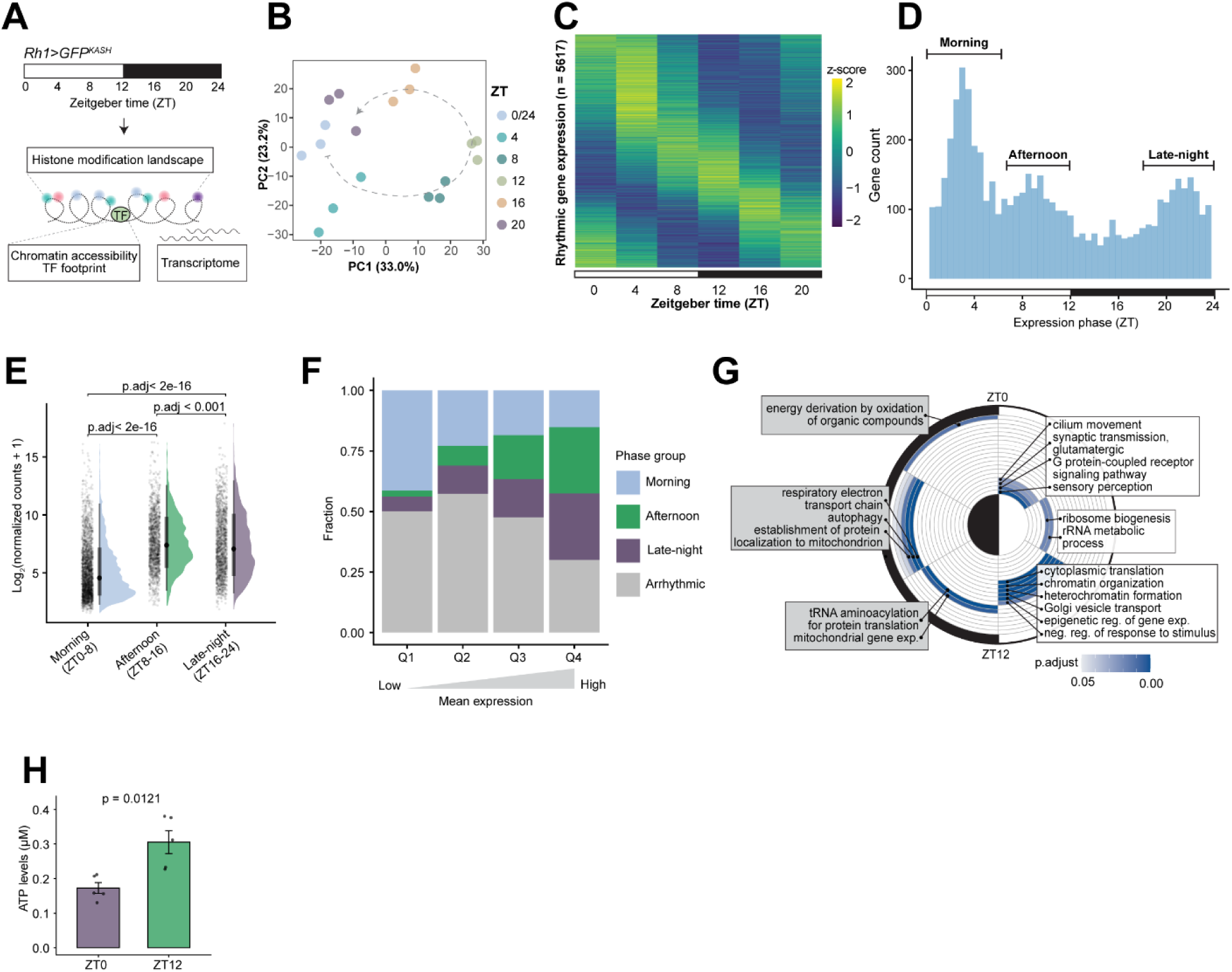
Rhythmic transcription is organized into waves across the diurnal cycle. (A) Photoreceptor nuclear RNA-seq, CUT&RUN and ATAC-seq in D10 male flies at 4-h intervals across a 24-h light:dark cycle (n = 3). (B) PCA plot of RNA-seq samples using regularized log transformed expression values. Arrow and dotted lines indicate the progression across ZT. (C) Heatmap of relative expression *z*-scores across ZT for rhythmic genes identified by dryR (p.adj ≤ 0.05). (D) Histogram of phase distribution for rhythmically expressed genes with morning, afternoon and late-night groups indicated by horizontal lines. (E) Raincloud plot of normalized expression levels across the three phase groups. Points represent genes, half-violin plots show expression level distribution with box plot indicating the median and interquartile range. Statistics were calculated using pairwise Wilcoxon test with Benjamini-Hochberg correction. (F) Bar plot showing the fraction of genes in each phase group across expression quartile. Q4 represents the highest expression quartile. (G) Representative biological processes gene ontology (GO) terms for rhythmically expressed genes across 4-h intervals. Terms are colored by adjusted *p* values. (H) Bar plot of ATP levels in D5 male fly heads at indicated ZT (n = 5). *P* value was calculated using student’s t-test.

Intriguingly, genes in these three phase groups show different average expression levels: the morning phase group showed significantly lower average expression than the afternoon or late-night groups (**Figure 1E**). Correspondingly, the most highly expressed genes most often peak in the afternoon or late-night (**Figure 1F**). As observed by other studies, highly expressed genes were also more likely to be rhythmic ^16,17^. Diurnal gene expression rhythms are thought to be important for both temporally separating incompatible biological processes and optimizing energy expenditure ^34,35^. Our data align with both theories: genes involved in cytoplasmic translation, one of the most energy-consuming processes in the cell ^36^, including ribosomal proteins and translation initiation factors, were enriched in the highly expressed afternoon group (**Figure 1G, Table S3**). In contrast, autophagy genes, whose activity is antagonistic to translation, were temporally separated showing peaks late at night, consistent with nighttime autophagy rhythms reported in *Drosophila* ^37^. Similarly, while genes involved in sensory perception and G-protein coupled receptor (GPCR) signaling pathway peak in the morning, genes involved in negative regulation of response to stimulus peak towards the end of the light phase. ATP levels in the avian retina oscillate across light:dark cycle ^6,38^, suggesting that production and consumption of energy may be tightly regulated across the diurnal cycle. Indeed, ATP levels in fly heads were significantly higher at the end of light phase (ZT12) than at its beginning (ZT0; **Figure 1H**). In *Drosophila* photoreceptors, because phototransduction is an energy-demanding process ^39^ and illumination substantially increases ATP cost ^40^, the timing of energetically costly gene expression programs, such as translation, may be constrained to times with higher ATP availability to balance biosynthetic and visual signaling demand. Together, these results suggest that daily transcription is temporally partitioned across the 24-h day according to both the relative expression level and biological function of each gene.

### The circadian clock transcription factors are insufficient to explain the phase of rhythmic gene expression

The molecular clock is regarded as the major regulator of circadian gene expression ^1^. However, in *Drosophila* photoreceptors, Clk and Cyc binding sites ^10^ only mapped to 232 unique genes (**Figure 2A, Table S4**), suggesting that Clk:Cyc do not directly regulate most of the 5617 rhythmic genes. When we clustered Clk:Cyc-bound genes into three groups (strong, moderate, weak) based on average Clk and Cyc CUT&RUN signals, only 68% of the Clk:Cyc-bound genes were classified as rhythmically expressed, regardless of the signal group (**Figure 2B**). Moreover, although Clk:Cyc occupancy is highly rhythmic, peaking at ZT12 and almost absent at ZT4, the phase distribution of Clk:Cyc-bound genes resembled all other genes, broadly peaking in the morning, afternoon, and late-night (**Figure 2A-B**). The discrepancy between the timing of circadian transcription factor binding and the phase of target gene expression has also been reported in mouse liver ^24,25^. We also noted that rhythmic Clk:Cyc-bound genes displayed a similar range of amplitudes and gene expression levels, except for the core clock genes that showed both the strongest Clk:Cyc signal and the highest expression amplitudes (**Figure 2C, Table S4**). Thus, while Clk and Cyc contribute to the likelihood of a gene being rhythmically expressed (**Figure S2D**), their binding alone is not sufficient to restrict the phase of their target expression to a specific time of day (i.e., ZT12.)

**Figure 2:**
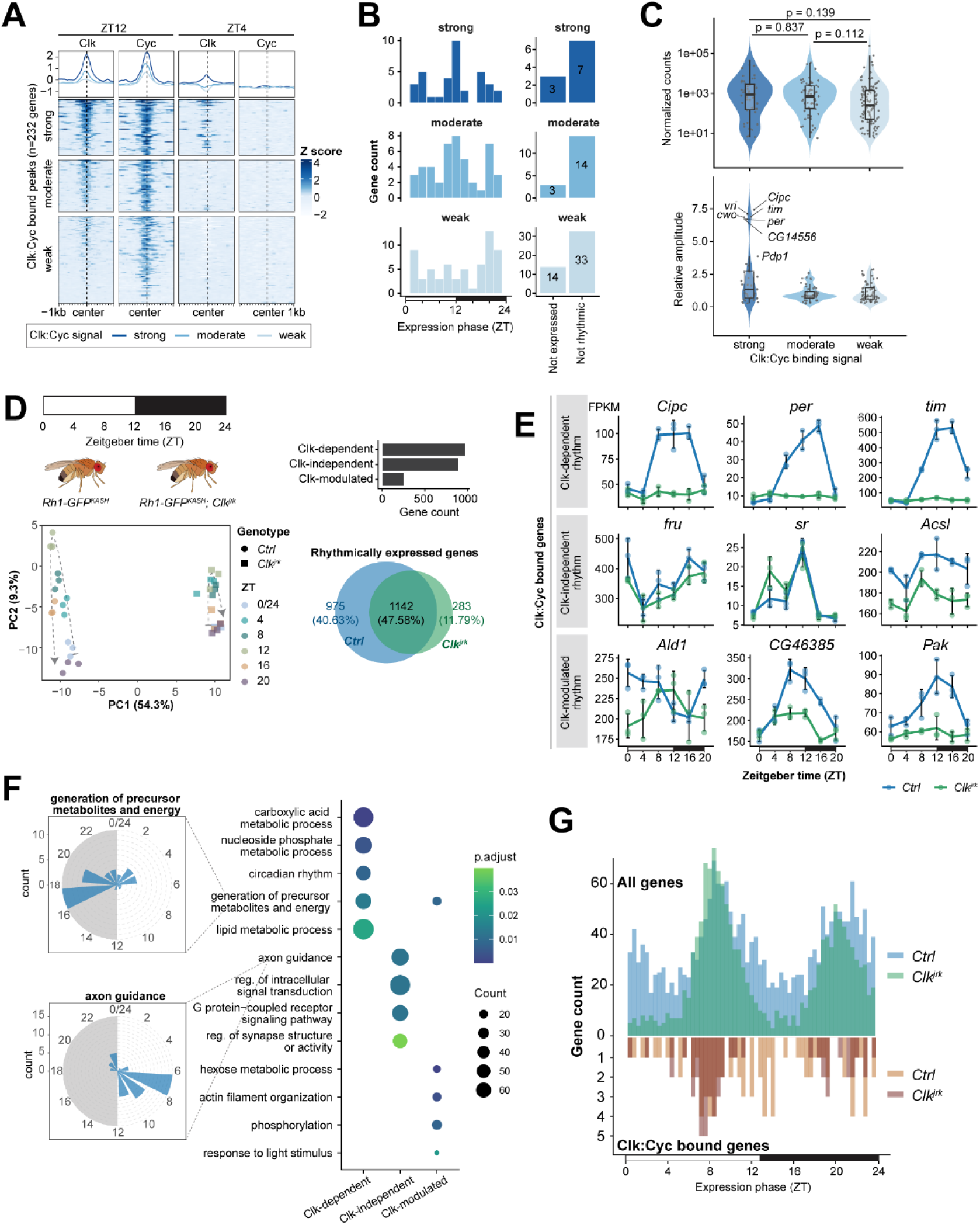
The circadian clock transcription factors are insufficient to explain the phase of rhythmic gene expression. (A) Gene metaplots and heatmaps of Clk and Cyc peak centers ±1 kb colored by relative signal (*z*-score) across the same region (rows) at ZT12 and ZT4 (n=3). Genes were stratified and colored by average Clk and Cyc signals. (B) Histogram of phase distribution for rhythmic Clk:Cyc-bound genes (left) and non-expressed and arrhythmically expressed genes (right) within each Clk:Cyc signal group. (C) Violin and box plots of normalized expression levels (top) and relative amplitude (bottom; log_2_-transformed peak-to-trough expression ratio) of genes in each Clk:Cyc signal group. *P* values were calculated using pairwise Wilcoxon signed-rank tests with Benjamini-Hochberg correction. (D) Photoreceptor nuclear RNA-seq from control and *Clk^jrk^* flies at 4-h intervals across ZT (n=3, top left). PCA plot of RNA-seq samples using regularized log transformed expression values (bottom left). Bar plot showing number of genes classified in each rhythm category in control and *Clk^jrk^* identified by dryR (top right). Venn diagram showing overlap of rhythmically expressed genes in control and *Clk^jrk^*(bottom right). (E) Line plots of representative Clk:Cyc-bound genes in each rhythm category. FPKM is shown as mean ± SD. (F) Representative GO terms in each rhythm category, colored by *p* values. Radial histogram showing phase of gene in selected GO terms. (G) Histogram showing phase distribution of all rhythmic genes (up, green and blue) and Clk:Cyc-bound genes (down, light and dark brown) in control and *Clk^jrk^*.

Next, we asked if circadian transcription factors were necessary for all diurnal transcription in photoreceptors. When we profiled the diurnal photoreceptor transcriptome of *Clk^jrk^* mutant flies ^41^ that are behaviorally arhythmic (**Figure S2A-B**), we found that only half of the genes that were rhythmic in control flies lost rhythmicity in the *Clk^jrk^* mutant (Clk-dependent genes, n = 975, **Figure 2D, Table S5**). These Clk-dependent genes include core clock genes, such as *Clock interacting protein circadian (Cipc)*, *period (per)*, and *timeless* (*tim)* (**Figure 2E**). However, other Clk:Cyc-bound genes, such as *fruitless* (*fru), stripe* (*sr)* and *Acyl-CoA synthetase long-chain* (*Acsl)*, either remained rhythmic in the *Clk^jrk^* mutant (Clk-independent, n = 890), or showed altered rhythmicity parameters such as phase or amplitude (Clk-modulated, n = 252, *Ald1*, *CG46385*, *Pak*). While Clk-dependent genes were involved in circadian rhythm and metabolism, the genes that remained rhythmic in the *Clk^jrk^* mutant were associated with sensory neuron function processes, such as signal transduction, axon guidance, and synapse organization (**Figure 2F, Table S6**). Intriguingly, we observed a compression in the overall phase of rhythmic transcription in the *Clk^jrk^* mutant, where the phases of rhythmically expressed genes were concentrated into two dominant time windows at ZT8 and ZT20 (**Figure 2G**). The same compression of phase windows occurs in mouse liver upon genetic disruption of the circadian clock, and has been termed *system-driven rhythms* ^42,43^. Overall, these results suggested, apart from the core clock genes, Clk and Cyc alone are insufficient to determine the phase of most of their direct target genes under light:dark conditions. Moreover, Clk appears to be dispensable for roughly half of rhythmically transcribed genes in photoreceptors, suggesting that light itself, or its absence, may contribute to transcription at the middle of the day and night.

### Transcription factor binding contributes but remains insufficient to determine the phase of rhythmic gene expression

Clk:Cyc are proposed to act combinatorially with other transcription factors – both by binding on shared targets, and by regulating the expression of downstream transcription factors ^9,44,45^. We therefore extended our analysis to other transcription factors by performing footprint analysis on our diurnal ATAC-seq data ^10^. Using TOBIAS ^46^, we characterized the temporal footprints of 220 expressed transcription factors based on their experimentally determined binding motifs from the CIS-BP database ^47,48^ (**Figure 3A**). Across accessible regions, the vast majority of transcription factors showed the highest activity at ZT12, the same time when chromatin shows highest accessibility at promoters ^10^. However, their putative rhythmic target genes were not expressed in a concerted manner at ZT12, but were broadly distributed into the same three phase groups observed previously (**Figure S3A-B**). Moreover, only 43.7% of rhythmically expressed genes had transcription factor binding events (**Figure 3B**). Thus, like Clk:Cyc, the inferred temporal binding of most transcription factors does not match the phase of their target gene expression, and many rhythmic genes do not in fact have transcription factor footprints.

**Figure 3:**
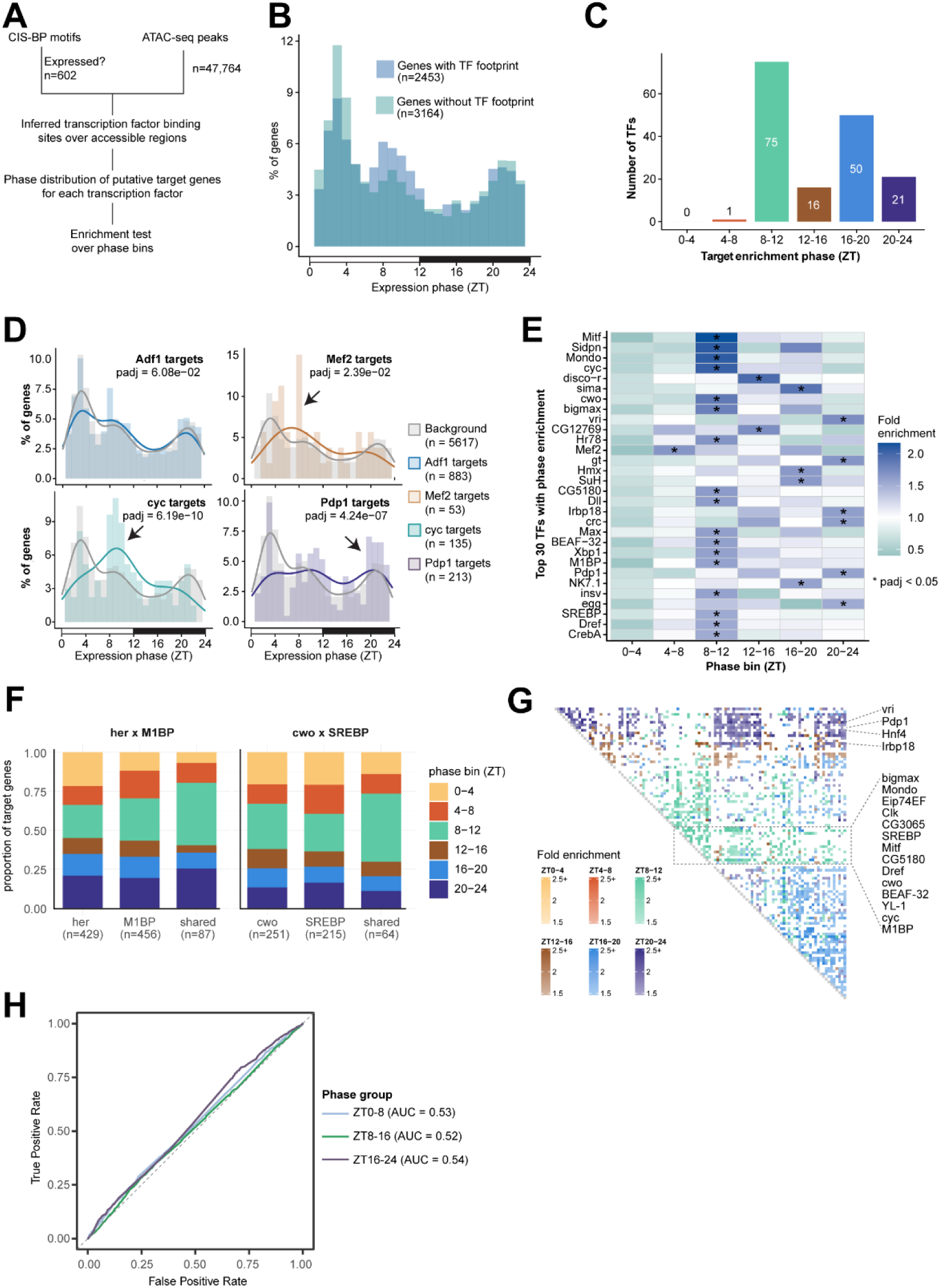
Combinatorial transcription factor binding contributes but remains insufficient to determine the phase of rhythmic gene expression. (A) Transcription factor footprint and target gene phase enrichment analysis based on diurnal ATAC-seq data using TOBIAS. (B) Phase distribution of rhythmic genes with and without transcription factor footprints. (C) Bar plot showing the number of transcription factors whose putative target genes are significantly enriched in indicated phase bin (p.adj ≤ 0.05). Putative target genes were defined as the genes nearest to each transcription factor’s bound footprints, annotated by TOBIAS. Phase enrichment was calculated as the proportion of a transcription factor’s putative target gene peaking at a given phase bin (ZT0-4, 4-8, 8-12, 12-16, 16-20, 20-24) relative to the corresponding background (all rhythmic genes) portion. Statistical significance was calculated using chi-square test with Benjamini-Hochberg correction. (D) Histogram with fitted density curves showing phase distribution of putative target genes for representative transcription factors versus all rhythmic genes (grey). Arrows indicate the phase bin driving enrichment. (E) Heatmap showing the top 30 transcription factors with the highest fold enrichment across each phase bin. Asterisks indicate phase bins with significant enrichment. (F) Bar plot showing proportion of target gene expression in each phase bin for individual transcription factors and their shared targets. (G) Heatmap showing fold enrichment of shared target genes for each transcription factor pair in the enriched phase bin; color indicates phase bin and intensity indicates fold enrichment. White space indicates no significant phase enrichment detected for given transcription factor pairs. The “phase-locking” transcription factors, showing enrichment on the same phase bin regardless of binding partners, were labeled on the right side. (H) Receiver operating characteristic (ROC) curves showing the performance of transcription factor footprint-based models in predicting the expression phase group of rhythmic genes.

We then wondered if, rather than dictating the phase for all their target genes, certain transcription factors may promote phase-specific transcription for a subset of their targets. Using a Chi-square enrichment test, we found that 163 transcription factors showed significantly different phase distributions for their target genes relative to all rhythmic genes (**Figure 3C, Table S7**). For example, Mef2 (Myocyte enhancer factor 2) targets were enriched at ZT4-8, cyc targets at ZT8-12, and Pdp1 (PAR-domain protein 1) targets at ZT20-24, while targets of other transcription factors, such as Adf1 (Adh transcription factor 1) showed no such enrichment (**Figure 3D**). Nevertheless, even phase-enriched transcription factors such as Pdp1 still bind many targets with broadly distributed expression phases, reminiscent of Clk:Cyc. In total, 75 transcription factors had targets enriched in the afternoon (phase bin ZT8-12, **Figure 3C**), including the clock transcription factors, Clk, cyc and cwo (clockwork orange); the architectural factors, BEAF-32 (Boundary element-associated factor of 32kD), Dref (DNA replication-related element factor), M1BP (Motif 1 Binding Protein) and CTCF; and Myc transcription factors, Mitf, Mondo, bigmax and Max (**Figure 3E**). 71 additional transcription factors had enriched targets in the late-night (ZT16-20 and ZT20-24), including the circadian transcription factors acting downstream of Clk:Cyc, vri (vrille) and Pdp1; and stress-responsive transcription factors, sima (similar), Irbp18 (Inverted repeat binding protein 18 kDa), crc (cryptocephal) (**Figure 3E**). In striking contrast, only a single neuron activity-dependent transcription factor, Mef2, had targets enriched among the morning genes (ZT4-8) and none was enriched between ZT0-4. These results suggest that transcription factors may favor the timing of a subset of their targets towards the afternoon or late-night, whereas morning genes may be governed by a distinct process.

Transcription factors often act cooperatively or antagonistically, and co-binding of Clk:Cyc with other transcription factors contributes to the transcriptional heterogeneity of Clk targets ^26,28,49^. We therefore asked whether co-binding by transcription factor pairs further restricts the expression phase of their shared targets beyond what either transcription factor achieves alone. We defined co-binding as inferred binding of two transcription factors at the same gene at the same ZT. Among the 163 transcription factors, shared targets of 2809 transcription factor pairs were phase-enriched relative to all rhythmic genes (**Table S8**). However, no pair showed stronger phase enrichment than the targets of either constituent transcription factor alone (**Table S9**). Although co-bound targets appeared to be more phase-enriched than individual target sets, such as genes co-bound by her (hermaphrodite) and M1BP tend to peak between ZT8-12 (**Figure 3F, Figure S3D**), this gain of phase enrichment may be largely due to the smaller size of co-bound targets. Instead, clustering of these co-binding transcription factors indicated that certain transcription factors constrained the phase of their target genes towards the same time window, regardless of their co-binding partner (**Figure 3G, Figure S3C**). These “phase-locking” transcription factors included Dref, M1BP, BEAF-32, Clk, cyc and cwo for the afternoon genes, and Pdp1, vri, Hnf4 (Hepatocyte nuclear factor 4), and Irbp18 for late-night genes. We next asked whether transcription factor binding predicts a gene’s phase group. Using multinomial logistic regression based on individual binding events of the 35 most robustly enriched transcription factors (p.adj ≤ 0.05, fold of enrichment > 1.5, with 5-fold cross validation), we observed poor predictive performance for all three phase groups (AUC ≈ 0.55) (**Figure 3H**). Thus, like Clk:Cyc, although some transcription factors favor afternoon or late-night transcription, the binding of these transcription factors is not sufficient to predict the phase of gene expression.

### Chromatin states segregate gene expression timing

Because the average expression levels differed significantly between the morning, afternoon, and late-night genes (**Figure 1E**), we considered if other genomic or epigenomic features might contribute to their phase of expression. HOMER motif analysis of promoters revealed that the morning-peaking genes were enriched for the Initiator (Inr) motif (**Figure 4A**), a simple core promoter element that is recognized by transcription factor IID (TFIID) and is sufficient for low level basal transcription ^50^. In contrast, afternoon-peaking genes were enriched for Dref and M1BP motifs that are associated with strong promoter activity and high expression levels ^51,52^, as well as E-box motifs bound by bHLH factors such as Clk, Cyc, and Myc ^53^. Importantly, the target genes of these architectural and E-box-binding transcription factors also preferentially peak in expression at ZT8-12, identified by phase enrichment analysis (**Figure 3E**). The late-night genes instead showed enrichment of the GAGA factor (Trl) motif, which is associated with RNA polymerase II (Pol II) pausing and chromatin accessibility ^54^. Thus, these phase groups differ in promoter architecture, suggesting that morning peaking genes rely more on Inr-associated basal transcription, while the afternoon and late-night genes are activated by sequence-specific transcription factors on more permissive chromatin.

**Figure 4:**
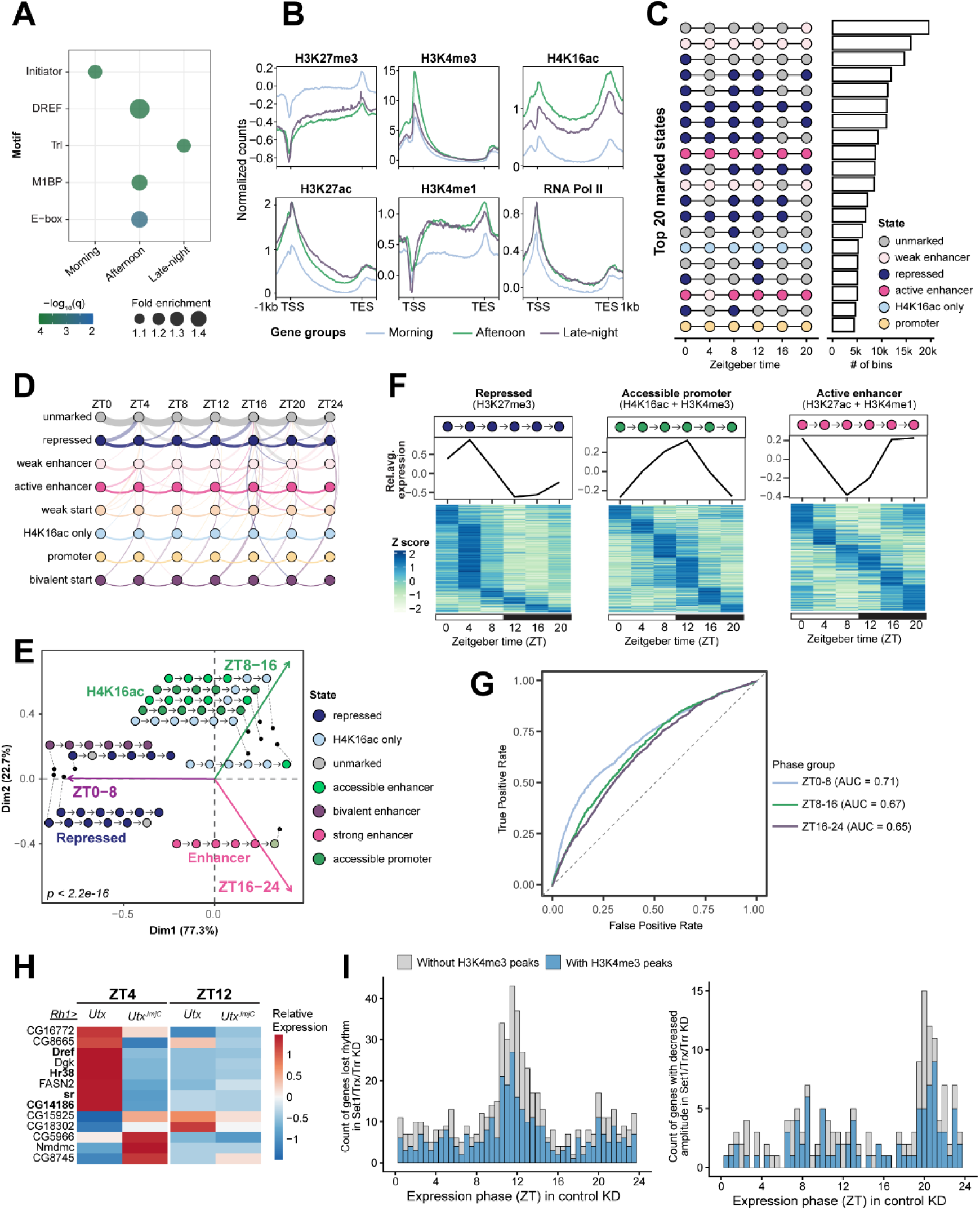
Morning peaking genes are responsive to changes in H3K27me3 while afternoon peaking genes are responsive to changes in H3K4 methylation. (A) Dot plot of promoter motif enrichment for each phase group from HOMER. Color represents statistical significance and size shows fold enrichment. (B) Gene metaplots showing mean normalized counts across all six ZTs for indicated histone marks in photoreceptors (n=3), separated by phase of gene expression, determined by nuclear RNA-seq. (C) Dot plot and bar plot of the top 20 most frequent chromatin states and number of 200-bp bins in these states. (D) Arc plot showing chromatin state transitions across 24 h. Dots represent chromatin states and the width of edges is proportional to the number of 200-bp bins that undergo transitions. (E) Biplot of correspondence analysis between chromatin states and gene expression phases. Colored arrows indicate three phase groups and dots represent best projected (cos^2^ > 0.5) chromatin states. (F) Heatmap showing expression *z*-scores and line plots of averaged relative expression for genes associated with indicated chromatin states. (G) Receiver operating characteristic (ROC) curves showing the performance of chromatin signature-based models in predicting the expression phase group of rhythmic genes. (H) Heatmap of relative expression of ZT-specific differentially expressed genes in photoreceptor-overexpression of *Utx* versus demethylase-dead control *Utx^JmjC^*, identified using a genotype × ZT interaction model from DESeq2. (I) Stacked histogram of phase distribution for rhythmic genes that show lost rhythm (left) or decreased amplitude (right) in all three H3K4 methyltransferase knockdowns. Phase values were acquired from mCherry control knockdown. The number of genes with and without H3K4me3 peaks were indicated by colors.

Histone modifications correlate with expression levels ^55^, and contribute to regulation of transcriptional cycle such as Pol II pausing and release ^56^. We previously found that genes with high levels of H3K4me2/3 tend to peak between ZT8-16, whereas genes with low levels peak between ZT0-4 ^10^. To extend this, we profiled H3K27ac and H4K16ac, as well as a repressive mark, H3K27me3, using CUT&RUN across the diurnal cycle in young photoreceptors. Building on the pattern suggested by H3K4me2/3, genes expressed at different phases displayed distinct epigenetic features: morning-peaking genes showed the highest overall levels of H3K27me3 and the lowest levels of all four active histone marks and RNA Pol II (**Figure 4B**). Conversely, both the afternoon and late-night genes carried relatively high levels of enhancer marks (H3K27ac and H3K4me1), with the afternoon genes showing substantially higher levels of the promoter marks H3K4me3 and H4K16ac (**Figure 4B**). We noted that while individual histone marks oscillated across the day, their oscillatory patterns were the same in all phase groups (**Figure S4A-B**), consistent with what we and other have previously observed ^10,27,57^. Thus, baseline chromatin environments, including both the type and levels of histone marks, rather than their oscillations, best distinguish rhythmic gene expression phases.

To further interrogate the association between chromatin features and transcription timing, we binned the genome into 200-bp segments and assigned each segment to one of 32 (2^5^=32) chromatin states defined by the presence of five histone marks ^58–60^, including H3K4me1, H3K4me3, H3K27ac, H4K16ac and H3K27me3 (**Table S10**). Most genomic segments stayed in the same chromatin state throughout the day or switched only between a given state and unmarked chromatin. The largest changes occurred in H3K27me3-marked repressive regions, many of which became unmarked from ZT0 to ZT4 (**Figure 4D, Figure S4C**). Using correspondence analysis, we compared the phase of expression and chromatin states for each rhythmic gene and identified a significant correlation (**Figure 4E**). H3K27me3 and bivalent states were associated with morning peaking genes (ZT0-8), with ∼ 70% of genes in H3K27me3-marked state peaking at ZT0-4. In comparison, regions marked by H4K16ac and H3K4me3 correlated with peak expression in the afternoon (ZT8-16), while those marked by H3K27ac and H3K4me1 modestly correlated with late-night genes (ZT16-24) (**Figure 4E-F**). We compared the enrichment of the phase-enriched transcription factors in each of these chromatin states and found that many of the afternoon and late-night transcription factors were preferentially enriched for specific chromatin states (**Figure S4D**). Matching HOMER analysis, M1BP and Dref from the afternoon transcription factor group were associated with promoter regions marked by H3K4me3 but were largely depleted at H3K27ac-marked regions, where instead the night transcription factor GAGA factor (Trl) was enriched (**Figure S4D**). Finally, we asked if these chromatin signatures could predict the phase of gene expression. For each gene, we stratified the levels of each histone mark into 4 groups (high, medium, low, and none) and used these as predictors in a multinomial logistic regression model of phase group. This chromatin-based model outperformed the transcription factor-based model (overall AUC = 0.68 vs. 0.55) and performed the best for the morning genes, suggesting this group may be more responsive to chromatin-based regulation (AUC = 0.71, **Figure 4G**). Together, these results show that chromatin signatures corresponding to modes of transcription regulation are associated with the phase of gene expression, suggesting that different chromatin environments promote specific diurnal timing of transcription.

### Morning peaking genes are responsive to changes in H3K27me3 while afternoon peaking genes are responsive to changes in H3K4 methylation

If chromatin signatures contribute to expression phase, then manipulating specific histone marks should alter gene expression in a phase-specific manner. To test this hypothesis, we focused on the H3K27me3 and H3K4me3 marks that showed strongest distinction across phase groups. Morning genes showed high levels of H3K27me3, low average expression levels, and enrichment of the Inr motif, suggesting they are basally transcribed in the morning and repressed at other time points. Since these genes are enriched for processes associated with sensory neuron function, we speculate that their expression might be rapidly induced in response to light onset. In developing neurons, activity-regulated genes often show a bipartite chromatin signature with H3K27ac-marked promoters and H3K27me3-marked gene body ^61^. While transcription is overall less productive on these bipartite genes, this chromatin signature enables rapid removal of H3K27me3 and stimulus-dependent gene induction ^61^. Notably, ∼ 3% of the genome was co-marked by H3K27me3 and either H3K27ac or H3K4me3 (**Figure S4C**), including the loci of activity-dependent transcription factors *Hr38 (Hormone receptor-like in 38,* ortholog of human *NR4A2)* and *sr (stripe,* ortholog of human *EGR1)* (**Figure S5**) ^62,63^. Therefore, we hypothesized that H3K27me3 sets the threshold for morning gene induction, and that reducing H3K27me3 would further enhance transcriptional induction. To test this, we overexpressed the H3K27me3 demethylase Utx or its catalytic inactive control Utx^JmjC^ in photoreceptors and performed nuclear RNA-seq at ZT4 and ZT12, corresponding to the peak and trough of morning gene expression. The main effect of ZT and genotype identified 247 and 133 differentially expressed genes, respectively. Only 11 genes showed a significant genotype and ZT interaction effect (**Figure 4H, Table S11**). These genes included activity-dependent genes *Hr38*, *sr*; *CG14186*, involved in Wnt signaling pathway and implicated in neuron activity ^64,65^; and the nutrient- and light-responsive genes, *FASN2 (Fatty acid synthase 2)*, and *Dgk (Diacyl glycerol kinase)*. These five genes showed higher expression only at ZT4, but not at ZT12 and carried a similar bipartite or repressive chromatin signature (**Figure S5**). Although the effect is modest, these results suggested that a subset of morning genes shares a similar regulatory mechanism with activity-dependent genes that enable their rapid transcriptional induction in response to stimulus.

Next, we examined the afternoon-peaking genes that are marked by high levels of H3K4me3 and enriched with promoter-associated transcription factors. To test whether H3K4 methylation is required for their afternoon expression, we turned to our previously published diurnal nuclear transcriptomes with individual Set1, Trx, or Trr knockdown across the 24-h light:dark cycle in young photoreceptors ^10^. Set1 is the major H3K4 tri-methyltransferase in larvae, and knockdown of either Trr or Trx also reduces H3K4me1/2/3 levels in adult photoreceptors ^10^. If H3K4me3 is necessary for rhythmic expression in the afternoon, then afternoon-peaking genes should be overrepresented among the genes that lose rhythmicity upon methyltransferase knockdown. Indeed, genes that peaked in the afternoon (ZT8-16) dominated the genes that lost rhythmicity (312 out of 576) in all three Set1/Trx/Trr knockdowns (**Figure 4I, Figure S4E**). Moreover, 58% of these 576 genes that lost rhythmicity were marked by H3K4me3 in the control knockdown. Notably, the H3K4 methyltransferase knockdowns did not affect all rhythmic genes equally: many genes with decreased amplitude – rather than completely lost rhythm – instead peaked preferentially in the late-night in these knockdowns (**Figure 4I**). These results suggested that afternoon genes are the most sensitive to the decrease of H3K4me3 and require all three methyltransferases for their rhythmic expression at around ZT12, whereas late-night peaking genes are affected but less sensitive to the knockdown of methyltransferases.

Overall, our data show that the phase of expression corresponds to chromatin states that are differentially bound by transcription factors. Afternoon genes were enriched with architectural, E-box transcription factors, and exhibited signatures of strong promoters, including high levels of H3K4me3, H4K16ac, and overall expression. Knockdown of H3K4 methyltransferases preferentially abolished rhythmicity of these afternoon genes. In contrast, genes peaking in the late-night had enhancer signatures, such as H3K27ac and H3K4me1, and were enriched for transcription factors that are either antagonistic or act downstream of the afternoon transcription factors. The morning genes differed from the other two groups and lacked phase-enriched transcription factors. Overexpression of Utx enhanced the morning expression of a subset of genes, including activity-dependent *Hr38* and *sr*, suggesting that relief of H3K27me3-mediated repression contributes to their morning induction. Together, we propose that these chromatin signatures restrict transcription to specific times of the day, at least in this light-responsive neuronal cell type.

## Discussion

Overall, our analysis suggests that the phase of diurnal transcription is shaped by a combination of chromatin signature and phase-enriched transcription factors, particularly the afternoon and late-night peaking genes. The insulator/architectural transcription factors enriched in the afternoon group, including M1BP, Dref, and BEAF-32, bind promoters and mediate long-range interactions that partition the genome into topologically associating domains (TADs) ^66–68^. Together with H3K4me3, they define TAD boundaries associated with high gene expression ^66–68^. We propose that these architectural transcription factors establish and/or maintain the TADs that segregate afternoon genes into a chromatin environment favoring high levels of transcription. E-box transcription factors ^69^, including Clk and Cyc, function within this environment, to promote phase-specific transcription of their targets towards the end of the light cycle. One way this compartmentalization could work is through biomolecular condensates, which concentrate the transcription machinery at target genes or sequester it away from non-targets ^70^. In mice liver, both BMAL1 and REV-ERBα form dynamic circadian condensates that are important for rhythmic gene expression ^71,72^, and in fly clock neurons, Period forms discrete puncta whose subnuclear localization, size and number oscillate over 24 h ^73^. The different levels of histone marks associated with the phase groups in our analysis may be important because histone marks can nucleate transcriptional condensates through multivalent interactions with the transcription machinery ^70^. For example, the local density of H3K4me3 and H3K27ac marks, measured by super-resolution microscopy, correlates with Pol II cluster sizes and transcriptional burst size ^74^. Thus, the chromatin states identified here may influence the spatial organization of the transcription machinery within the nucleus, thereby contributing to the temporal partitioning of gene expression across the day.

Late-night-peaking genes are distinguished from the afternoon group by both their enhancer-like chromatin signatures and transcription factor networks. Many late-night transcription factors are themselves transcriptionally controlled by afternoon-enriched transcription factors. For example, Pdp1 and vri, enriched in late-night genes, are direct targets of Clk:Cyc that require Clk activity for rhythmicity ^75^, and their mammalian counterparts REV-ERBα and NFIL3 have been implicated as master regulators of late-night genes in mouse liver ^30^. Other late-night transcription factors are known to antagonize or be suppressed by the function of afternoon-enriched transcription factors, suggesting that their opposing roles could lead to a switch between transcription of the afternoon and late-night genes. For example, BEAF-32 binding has been postulated to weaken pausing induced by the late-night transcription factor Trl ^76,77^, and M1BP not only promotes ribosomal gene transcription ^78^, but also suppresses expression of autophagy and apoptosis genes ^79^, which typically peak at late-night. Because late-night genes showed enhancer-like chromatin signatures, we propose that they depend more on transcription factor activity at enhancers. Consistent with this, enhancer RNA analysis in mouse liver indicates that enhancers were most active at late-night, while transcriptional regulation in the afternoon seems to occur primarily at promoters ^29,30^. It is unclear whether the chromatin signatures we observe in fly photoreceptors are also found at rhythmic genes in mouse liver, but the parallel association of enhancer- and promoter-based regulatory mechanisms with distinct phase groups suggests that a combinatorial model of chromatin signatures and transcription factor binding may be a conserved mechanism to temporally regulate gene expression in peripheral clock tissues.

The morning genes differed from the other two groups and seem to be regulated predominantly via chromatin state rather than transcription factors. In neurons, activity evoked by external stimulation or spontaneous firing induces the expression of activity-dependent genes, many of which encode transcription factors that activate downstream gene expression important for synapse formation and function ^80^. In *Drosophila* photoreceptors, light activates a G-protein coupled receptor phosphoinositide signaling cascade that leads to influx of cations such as calcium, and depolarization of the neuron ^81^. Thus, light onset at ZT0 would be expected to induce the expression of activity-dependent genes, and indeed, the immediate early transcription factors *Hr38* and *sr* are present in the morning group. Notably, *Hr38* or *sr* overexpression expands the projection of small ventrolateral neuron (s-LNv), which is circadian regulated occurring in the early morning ^63^. Our data indicated that H3K27me3 normally keeps morning gene expression at low levels and restricts their induction to a specific time window, a mode of regulation similar to the induction of activity-dependent genes in developing neurons that carry bipartite chromatin signature ^61^. Reduction of H3K27me3, whether via its transient dip at ZT4 or by Utx overexpression, may lower the threshold for inducing this morning wave of transcription. Indeed, in cultured cells, degron-induced depletion of the PRC1/2 complexes leads to rapid induction of H3K27me3-marked genes, with lowly expressed ones showing the earliest transcriptional response within 2h of depletion ^82^. Thus, unlike afternoon and late-night genes, morning genes may be gated mainly by chromatin state and induced by environmental stimulus.

The transcription factor networks associated with specific phases in photoreceptors include both core clock regulators and many other transcription factors. While critical for maintaining rhythmic behavior and gene expression in the absence of zeitgebers ^83^, the circadian clock may have a more nuanced role in diurnal rhythmic gene expression, particularly in the retina. We found that the transcriptional waves at ZT8 and ZT20 are largely maintained in the *Clk^jrk^* mutant. In the mouse liver, similar *system-driven genes* remained rhythmic at comparable times in both *Bmal1* and *Cry1/2* knockouts, which has been postulated to be regulated via signals from the SCN ^42^. However, because rhythms in the pacemaker neurons are also disrupted in the *Clk^jrk^* mutant ^84^, system-driven rhythms are unlikely to depend on the central clock and may be driven by external stimulus. In mouse liver, mTOR signaling is both necessary and sufficient to mediate hepatic system-driven rhythmic gene expression in the absence of a functional clock ^43^, suggesting that phosphorylation may represent a critical signaling mechanism through which zeitgebers exert their temporal control on transcription. While nutrient levels are a crucial zeitgeber in the liver, light has a more critical role in the retina ^4,6^. Indeed, light exposure increase phosphorylation of more than 1000 proteins in *Drosophila* heads ^85^ and many phosphorylation sites show circadian rhythms ^86^. In the mouse forebrain, around half of phosphoproteins exhibit rhythmic phosphorylation, nearly all of which is abolished by sleep deprivation ^87^. These diurnal phosphorylation events may provide clues to the transcriptional mechanisms that drive light-dependent expression rhythms, independent of the molecular clock.

We propose that regulating diurnal gene expression rhythms through a combination of chromatin signature and transcription factors enables both the temporal segregation of biological processes and optimization of transcriptional timing based on energy availability. Photoreceptors are among the most energetically demanding cells: visual responses including active ion transport, maintenance of ionic gradients, and synaptic transmission consume vast amount of ATP ^88,89^. Therefore, timing high levels of transcription, and the translation that follows, to time windows of lower phototransduction demands ^40^ could reduce the competition for metabolic resources such as ATP and GTP ^90^, especially for highly expressed genes which impose greater energy demand. Consistent with this possibility, afternoon and late-night genes were expressed at higher levels than morning genes, and ATP is more abundant at the end of the light phase at ZT12 compared with early morning at ZT0 (**Figure 1H**). In addition, translation in *Drosophila* clock cells peaks at CT9 and CT20 ^91^, which overlaps with the latter two transcriptional waves observed in this study, suggesting global transcription and translation are temporally synchronized. As in the mammalian retina, *Drosophila* photoreceptors are long-lived neurons that maintain their function over the lifespan of the organism, up to 80 days in flies ^92^. The chromatin signatures associated with the afternoon and late-night genes may both promote their diurnal expression and help maintain these rhythms over time. Supporting this idea, H3K4me3 has been proposed to promote transcription consistency by retaining memory about prior transcriptional status ^93^. We propose that these chromatin signatures provide long-term and efficient regulatory mechanisms that collaborate with transcription factors to maintain diurnal rhythms and optimize photoreceptor function across the light cycle.

## Methods

### Fly stocks and genotype

*Drosophila* were maintained on standard cornmeal food at 25°C under 12-h light:12-h dark conditions. Male flies were collected on day 10 ± 1 day post eclosion at indicated ZT ± 15 min with dark phase collections (ZT0, ZT16 and ZT20) performed under dim red light. *Rh1-Gal4>UAS-GFP^KASH^* flies were collected for CUT&RUN. *Rh1-GFP^KASH^; Clk^jrk^* was generated in this study using parental BDSC stock (RRID:BDSC_80927) and used for photoreceptor nuclear RNA-seq. For overexpression RNA-seq, *Rh1-Gal4>UAS-GFP^KASH^* were crossed to either *UAS-Utx* or control *UAS-Utx^JmjC^* (RRID:BDSC_94164, BDSC_94165) and F_1_ male progeny collected on day 10 ± 1 day at ZT4 or ZT12. A full description of all stocks used in this study is provided in Table S1.

### Circadian locomotor activity assay

Flies were raised in 12-h light:12-h dark conditions and males were transferred to vials for locomotor activity assays at 5 days post-eclosion. Locomotor activity was monitored using the DAM2 *Drosophila* activity monitor (Trikinetics) for 5 days of 12-h light:12-h dark followed by 5 days of constant darkness. Data analysis was performed using ShinyR-DAM ^94^.

### Nuclei immuno-enrichment (NIE)-based RNA-seq and CUT&RUN

Fly heads were isolated, homogenized and then incubated with anti-GFP coated magnetic beads to immuno-enrich for GFP-tagged photoreceptor nuclei. Bead-bound nuclei were then used for RNA-seq and CUT&RUN as described previously ^10,95,96^. Detailed protocols is available at ^97^.

#### RNA-seq

nuclear RNA extraction and purification was performed using Zymo Direct-zol RNA Microprep Kits (Cat. #R2060). rRNA was depleted using customized *D. melanogaster* rRNA probes and NEBNext® RNA Depletion Core Reagent Set (Cat. #7865). cDNA library was constructed using NEBNext® Ultra™ II Directional RNA Library Prep Kit for Illumina® (Cat. #7760). Pair-end (PE150) sequencing was performed on AVITI24 platform with Genomic Center of University of Minesota.

#### CUT&RUN

bead-bound nuclei from the same biological sample was split evenly for different histone antibodies. The following antibodies were used: anti-IgG (EpiCypher, 13-0042, RRID: AB_2923178); anti-H3K27ac (Abcam, ab4729, RRID: AB_2118291); anti-H4K16ac (Abcam, ab109463, RRID: AB_10858987), anti-H3K27me3 (EpiCypher, 13-0055, RRID: AB_3665059). CUT&RUN libraries were constructed using NEBNext® Ultra™ II DNA Library Prep Kit for Illumina® (Cat. # E7645L). Pair-end (PE150) sequencing was performed on NovaSeq X platform with Medical Genomics Core of Indiana University.

### RNA-seq, CUT&RUN, ATAC-seq data analysis

Data processing was as described in ^10^. Briefly, adaptor sequences and low-quality reads were trimmed using trimmomatic ^98^. Trimmed reads were aligned to the *D. melanogaster* genome (BDGP6.54) using HISAT2 ^99^ (RNA-seq) or Bowtie2 ^100^ (CUT&RUN). SAM, BAM and bigWig files were generated using SAMtools and deepTools ^101,102^. Quantification of RNA-seq samples were performed using featureCounts for exons ^103^. Gene expression rhythmicity analysis was performed using dryR (p.adj ≤ 0.05 for D10 nuclear RNA-seq; BICW ≥ 0.6 for *Clk^jrk^* RNA-seq) ^42^. Differential gene expression analysis was performed using DESeq2 ^104^.

#### Transcription factor footprinting analysis

Transcription factor DNA binding motifs were acquired from the CIS-BP database (v. 3.00) ^105^ for *D. melanogaster.* A total of 602 DNA binding motifs (corresponds to 220 transcription factors) were used as input for footprint analysis after two steps of filtration: transcription factor expression level (CPM ≥1 in all three biological replicates in at least one timepoint) and motif redundancy (removed if motif position weight matrices similarity > 99.9% using PWMSimilarity function in the TFBSTools R package ^59,106^). A master peak list was generated for all accessible regions defined by ATAC-seq data and used for footprinting. Briefly, peaks were called using MACS2 with FDR < 0.01 ^107^ and peaks present in more than two biological replicates in the same condition were retained using BEDTools ^108^. Peaks within 120bp were then merged from all six timepoints to generate the master peak list. Footprinting was analyzed using TOBIAS ^46^, which corrects for inherent Tn5 insertion bias (ATACorrect), predicts transcription factor binding by accounting for local accessibility (ScoreBigwig), and detects differential transcription factor binding across conditions (BINDetect), and annotate footprints to nearest genes ^46^. A customized blacklist for *D. melanogaster* ^10^ was applied to minimize bias estimation for footprinting.

#### Transcription factor target gene phase enrichment analysis

Transcription factor target genes were defined from TOBIAS footprints restricted to regulatory regions within promoters and first introns. Target gene phase enrichment was determined by Chi-square goodness-of-fit test comparing the phase distribution of each transcription factor’s putative targets to the background distribution of all rhythmically expressed genes. Phases of both background gene and target gene sets were binned into six 4-h intervals (ZT0-4, ZT4-8, ZT8-12, ZT12-16, ZT16-20, ZT20-24). The fold of enrichment was calculated as the proportion of target genes in each phase bin relative to the corresponding background portion. *P* values were adjusted for multiple testing using the Benjamini-Hochberg, and transcription factors with a p.adj ≤ 0.05 were considered significantly phase enriched.

#### Transcription factor co-binding and network analysis

Co-binding and network analysis were performed for transcription factors with significant phase enrichment (n = 163, p.adj ≤ 0.05). Two candidate transcription factors were defined as co-binding when their footprints fell within the same ATAC peak at the same timepoint. Phase enrichment analysis for shared target genes of each transcription factor pair was performed similarly using Chi-square test as described above. Enrichment from 1395 transcription factor pairs were significant out of 5468 total pairs compared with all rhythmic gene background (Table S8, transcription factor with less than 20 co-binding partners were dropped for co-binding matrix). Nodes represent transcription factors and edges represent pairs whose shared target genes show significant phase-specific enrichment (chi-square test, p.adj ≤ 0.01, fold of enrichment ≥ 2, number of shared target genes ≥ 15). Node size represented each transcription factor’s total footprint count; node color represented the dominant phase its targets were enriched in. Layout was computed using a stress-minimization algorithm and transcription factors were grouped to improve visual separation of related clusters using igraph ^109^.

#### Chromatin state analysis

Combinatorial chromatin states across the 24-h day were assigned genomic regions using the chromstaR package ^58,60,110^. The genome was partitioned into 200-bp, non-overlapping bins for multivariate peak calling for H3K4me1, H3K4me3, H3K27ac, H4K16ac and H3K27me3. Based on the presence/absence of each mark at each ZT, a discrete chromatin state is assigned to every genomic region, acquiring 2^(5 histone marks x 6 timepoints) total possible states (mode=”differential”). Two samples (ZT08_H3K27ac_rep2 and ZT16_H3K27me3_rep1) were not included in this analysis due to poor correlation with their corresponding biological replicates, which would negatively impact joint peak calling. The posterior probability for the Hidden Markov Model used by chromstaR were set at 0.9999 to adjust the strictness for histone mark peak calling. Only the most abundant chromatin states and standard chromosomes (2L, 2R, 3L, 3R, 4, X, Y) were considered for downstream analysis.

Genomic regions in each state were then annotated to their nearest genes using ChIPpeakAnno package ^111^. The relationship between chromatin states and rhythmical gene expression was extracted using correspondence analysis and visualized on a biplot with the top contributing chromatin states with cos^2^ > 0.5 shown (**Figure 4D**), using Factoextra and FactoMinerR packages ^112^. Statistical significance for this association was determined using a Chi-square test.

To assess transcription factor binding preferences over chromatin states, transcription factor footprints identified by TOBIAS were intersected with chromatin state segments. For each transcription factor, footprint enrichment in each state was calculated as the number of footprints normalized to the total footprints for that transcription factor, divided by the genomic coverage of the corresponding chromatin state, accounting for size differences in transcription factor binding sites and chromatin states. Statistical significance was calculated by fisher’s exact test followed with Benjamini-Hochberg correction.

#### Homer motif analysis

Motif analysis was performed for rhythmic genes in each phase group (ZT0-8, ZT8-16, ZT16-24) using HOMER (v. 4.11, Hypergeometric Optimization of Motif EnRichment) ^113^. Enrichment analysis for promoter regions of target genes in each phase group was performed relative to a matched background rhythmic gene set using *D. melanogaster* BDGP6.54 genome assembly as reference. Known-motif comparison was restricted to the insect motif database.

#### Multinomial regression analysis

Gene’s expression phase was modeled as a function of chromatin features and transcription factor footprints using penalized multinomial logistic regression, comparing candidate predictor sets under nested cross-validation, with model settings tuned only on the training portion of each split. For chromatin-based model, average levels for each histone mark across six ZTs were quantified using multiBigWigSummary in deepTools ^102^ and discretized to ordinal factors (none/low/moderate/high) as predictors. For transcription factor-based model, the top 35 transcription factors with most robust target phase enrichment (p.adj ≤ 0.05, fold of enrichment > 1.5) were included and their footprints across ZTs were collapsed to dummy variables of 0 (absent) and 1 (present) as predictors. Gene expression phases were binned into three groups (ZT0-8, ZT8-16 and ZT16-24) and used as outcome variables. Model performance was summarized as macro-averaged area under the receiver operating characteristic curve (AUC), computed from out-of-fold predicted probabilities pooled across the five-fold cross validation, using pROC ^114^; per-phase-group AUCs were reported separately.

### ATP measurement

ATP measurements were conducted using the ThermoFisher ATP Determination Kit (Cat. #A22066). 100 male flies (*w^1118^*) were collected on day 5 ± 1 day post eclosion at ZT0 and ZT12 (n=5). Fly heads were homogenized in cell lysis buffer (stock: 200mM Tris-HCl pH 7.5, 2M NaCl, 20mM EDTA, 0.2% Triton X-100) with Dounce homogenization on ice. Homogenates were centrifuged at 16,000xg for 1 minute and kept on ice. Clarified cell lysate was added to the reaction solution recipe provided in the kit in a ratio of 1:10. The reaction was incubated for 2.5 minutes at room temperature and luminescence was measured with a CLARIOstar (BMG Labtech). ATP values were determined from an ATP standard curve developed in tandem with sample measurements. Significant results were determined with student’s t-test.

### Data visualization

Gene ontology (GO) analysis was performed using clusterProfiler ^115^ and visualized using ggplot2 ^116^. CUT&RUN data were CPM-normalized and subtracted for respective IgG controls, and visualized with heatmaps and metaplots using deepTools ^102^. Genome track visualization for selected genes was plotted using the Gviz package ^117^.

## Date availability

High-throughput sequencing data is available at the Gene Expression Omnibus (GEO). *Data generated in this study:* GSE304767: photoreceptor nuclear RNA-seq for *Clk^jrk^* and control across 24-h diurnal cycle (36 samples); GSE317820: CUT&RUN for D10 H3K27ac, H4K16ac, H3K27me3 and IgG (288 samples); GSE349673: photoreceptor nuclear RNA-seq for Utx overexpression or Utx^JmjC^ at ZT4 and ZT12. *Previously published data analyzed in this study:* GSE262862: D10 photoreceptor nuclear RNA-seq; GSE253177: Photoreceptor Clk and Cyc CUT&RUN; GSE278041: D10 and D50 ATAC-seq, RNA Pol II, H3K4me1/2/3 CUT&RUN; GSE262534: photoreceptor mCherry RNAi, Set1 RNAi, Trx RNAi, Trr RNAi nuclear RNA-seq. Photoreceptor nuclear RNA-seq data for rhythmic gene expression in *Clk^jrk^*can be viewed here: https://gaoyameng.shinyapps.io/clkjrk_shinyr/

## Author contribution

G.M. generated CUT&RUN data for H3K27ac, H4K16ac and H3K27me3, analyzed the data, conceptualized the work, prepared figures, wrote and edited the manuscript. S.E.M generated *Clk^jrk^* RNA-seq data, performed circadian behavior experiments. L.D.K performed measurements of ATP. R.I.M.-Z. and G.M. generated data analysis pipeline. V.M.W conceptualized the work, supervised this study, designed experiments, wrote and edited the manuscript.

## Acknowledgements

Research reported in this publication was supported by the National Eye Institute of the National Institutes of Health under Award Number R01EY033734 to V.M.W and National Science Foundation under Award Number 2409483 to V.M.W. The content is solely the responsibility of the authors and does not necessarily represent the official views of the National Institutes of Health. Additional support was provided by Bird Stair and Howard Zalkin Award from the Purdue University Department of Biochemistry to G.M.

## Declaration of generative AI and AI-assisted technologies in the manuscript preparation process

During the preparation of this work, G.M. used Claude for providing coding suggestions for data analysis, and for checking grammar and language refinement. G.M. and V.M.W. reviewed and edited the output as needed and take full responsibility for the content of the published article.

---

**Figure S1:**
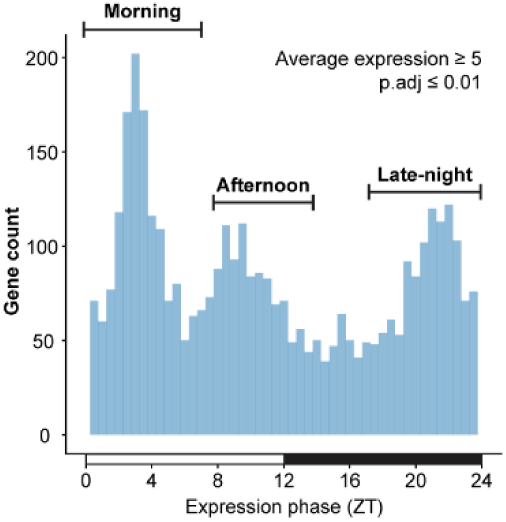
Similar triphasic distribution is observed with more stringent expression and rhythmicity cutoffs. Histogram showing phase distribution of rhythmic genes with log_2_-normalized counts ≥ 5 and p.adj ≤ 0.01, related to Figure 1A (CPM ≥1 in all three biological replicates in at least one ZT, p.adj ≤ 0.05)

**Figure S2:**
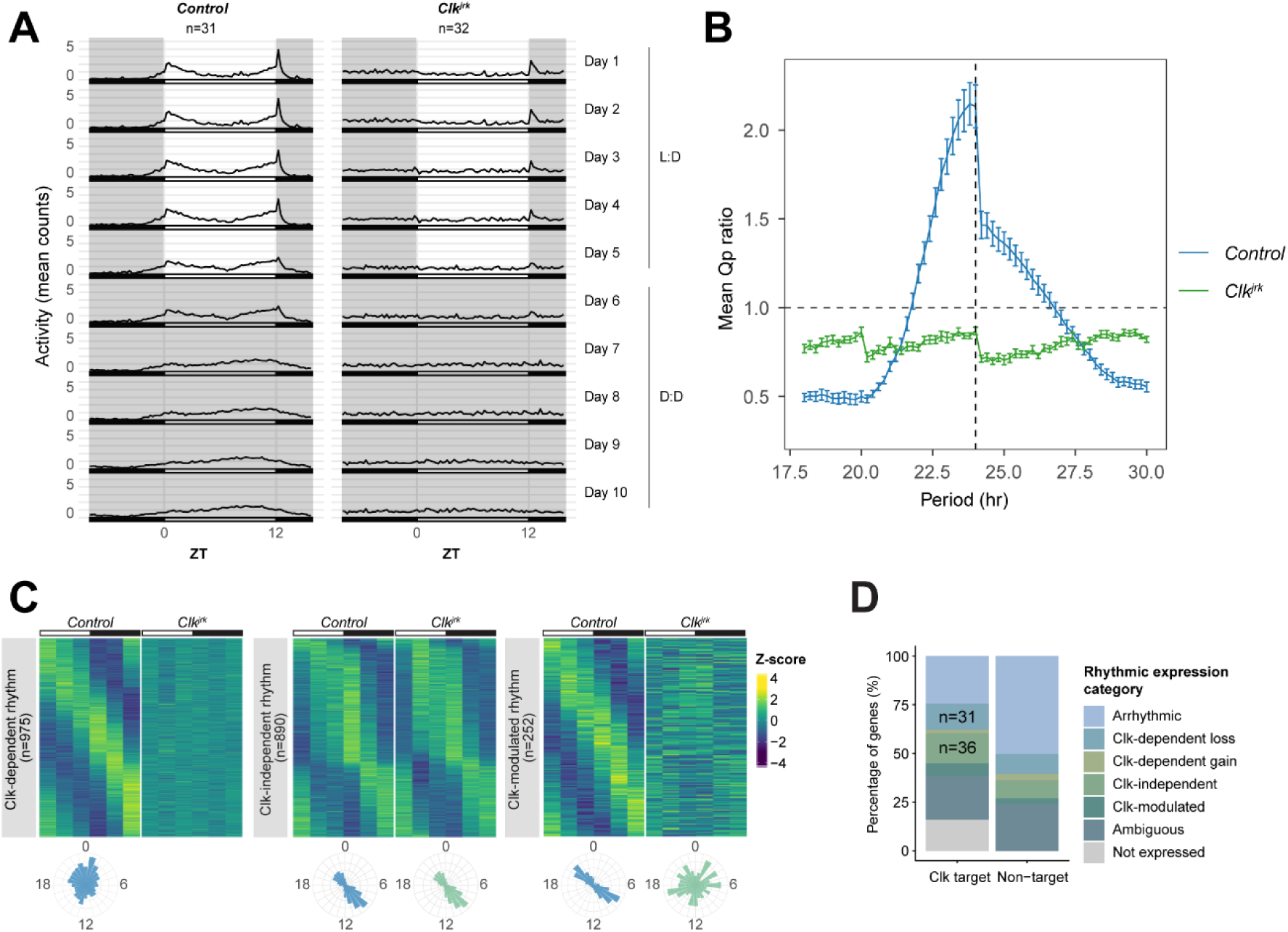
*Clk^jrk^*mutant is behaviorally arrhythmic and disrupts rhythmicity of approximately half of rhythmic genes in photoreceptors. (A) Locomotor activity actograms of mean counts of each genotype (*n* indicated) in 15-minute bins for 5 days of 12-hr:12hr L:D followed by 5 days of complete darkness (D:D). (B) Chi-square periodogram analysis of locomotor activity rhythms. The ratio of observed to significance threshold periodogram power (Q_P_ ratio = Q_act_/Q_sig_) is plotted as a function of tested period. Each point on the curve represents the mean Q_act_/Q_sig_ value for a given genotype, with error bars indicating the standard error of the mean (SEM). The horizontal dashed line denotes the threshold for significant rhythmicity, and the vertical dashed line marks a 24-hr period. (C) Heatmap showing *z*-scores of relative expression and radial histograms showing phase of genes in each rhythmic category defined by dryR. (D) Bar plot showing percentage of genes in each rhythmic category for Clk:Cyc-bound genes (targets) and non-Clk:Cyc targets. “Ambiguous” refers to genes with BICW ≤ 0.6.

**Figure S3:**
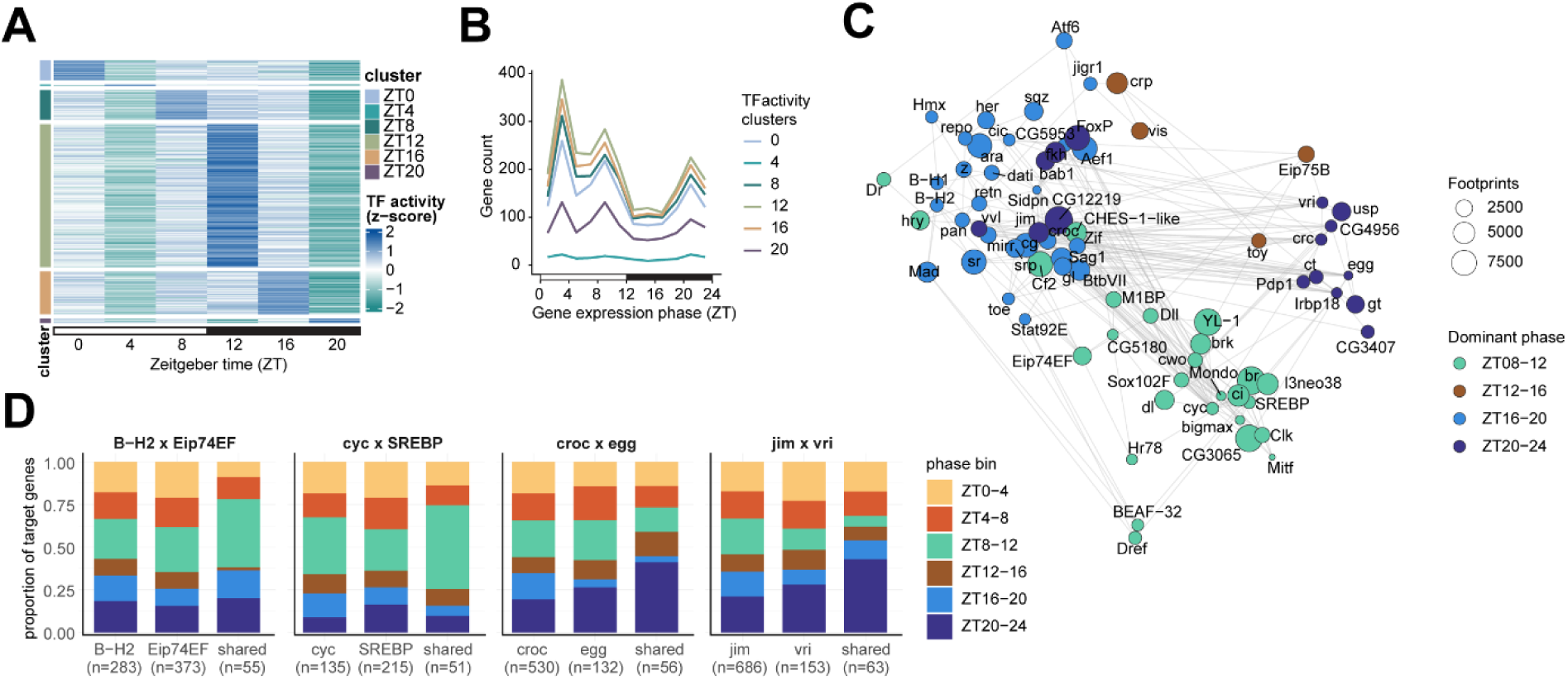
Transcription factor activity does not correlate with expression phase, but individual and combination of transcription factors show phase enrichment. (A) Heatmap showing clusters of transcription factor activity across ZT, grouped by the timing of peak transcription factor activity. Each row represents a motif, and activity scores were calculated across accessible regions using TOBIAS. (B) Line plot showing the phase distribution of putative target genes bound by transcription factors in each cluster. (C) Network showing transcription factor co-binding relationships. Nodes represent transcription factors and colored by the phase bin their putative target genes were enriched in. Edges represent transcription factor pairs whose targets show significant phase enrichment, and the sizes of nodes represent number of footprints given transcription factor.

**Figure S4:**
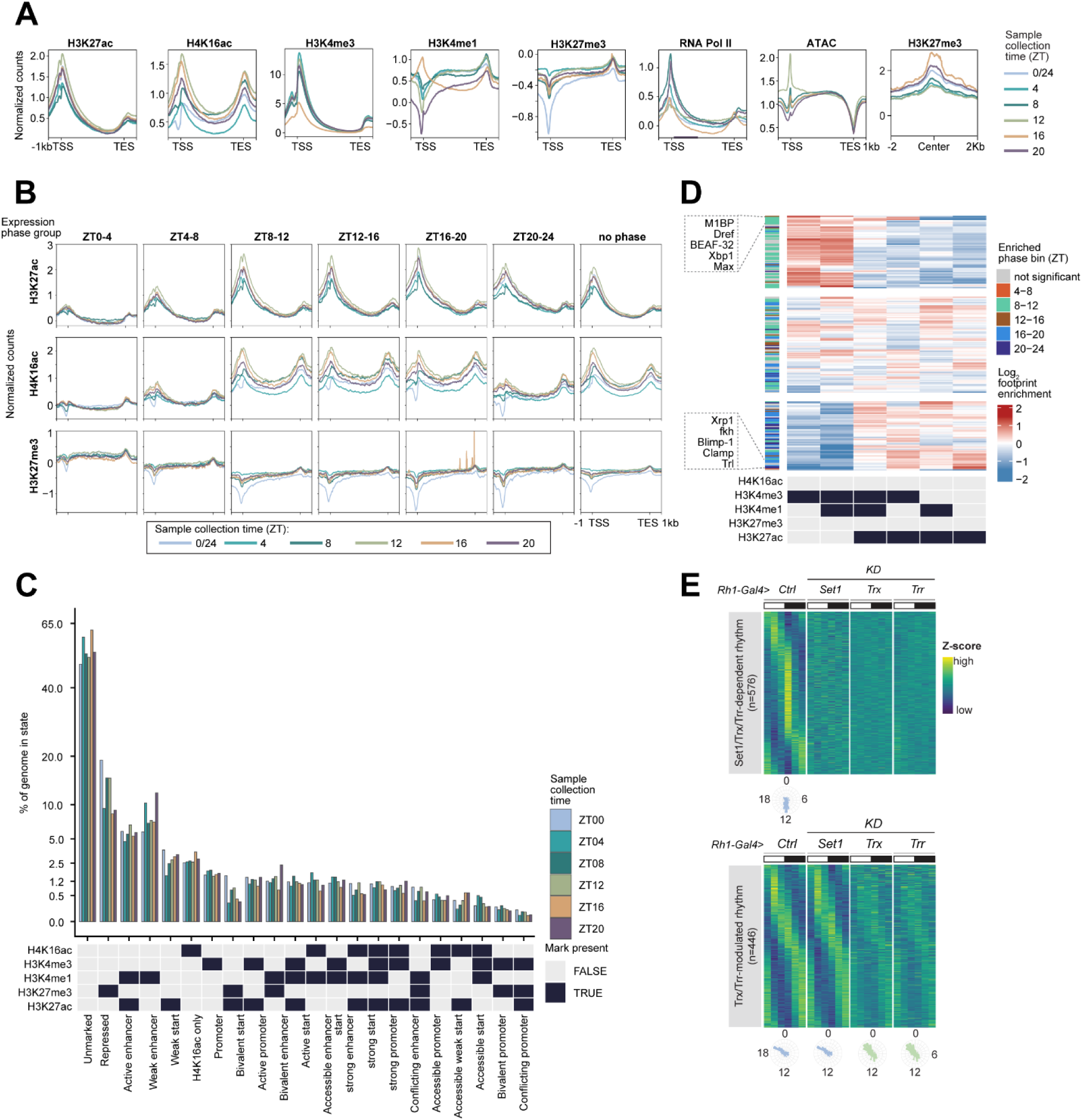
Temporal oscillation of histone marks across the day does not correlate with the phase of gene expression, which is instead correlated with chromatin states. (A) Gene metaplots showing mean normalized counts for RNA Pol II, chromatin accessibility and indicated histone marks in photoreceptors (n=3) at each ZT across 24-h day. (B) Gene metaplots showing mean normalized counts for indicated histone marks in photoreceptors for each phase group at each ZT across 24-h day. (C) Histogram showing the percentage of genome enriched in the indicated chromatin states at each ZT across 24-h day. Colored boxes indicate if histone mark is present in the state. (D) Heatmap showing fold enrichment of transcription factor footprints over indicated chromatin states, normalized to total footprints and chromatin state genomic coverage. Each row represents a unique transcription factor, colored by phase bin that their putative target genes were enriched in. Representative transcription factors were labeled. (E) Heatmaps showing expression *z*-scores for genes identified in each rhythmic category in mCherry (Ctrl), Set1, Trx and Trr knockdowns, and radial histograms showing phase of peak gene expression.

**Figure S5:**
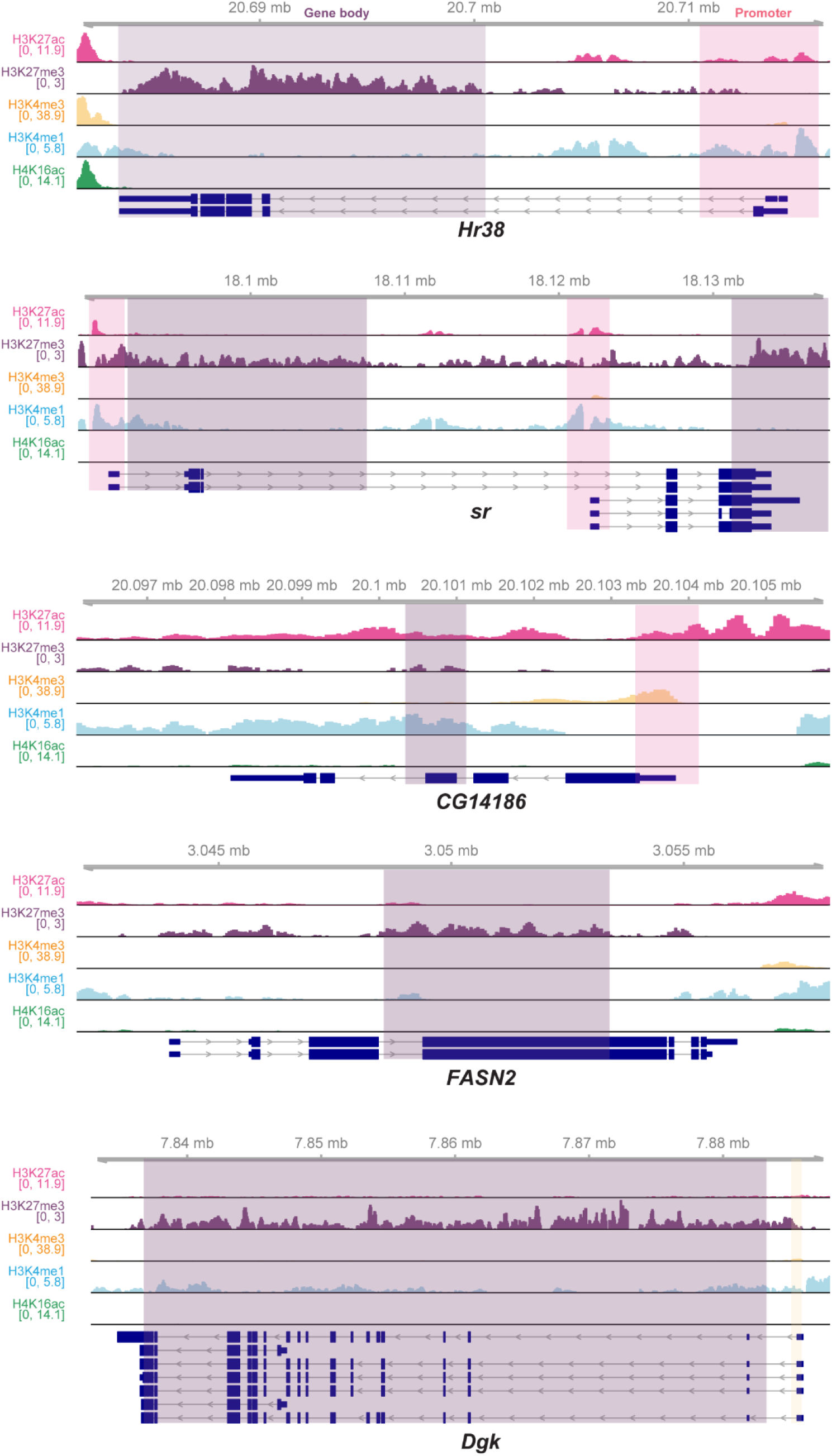
Activity-dependent genes exhibit bipartite or repressive states. IgG-normalized CUT&RUN genome browser profile of representative genes with bipartite (*Hr38, sr, CG14186*), bivalent (*CG14186*), and repressive features in young (D10) fly photoreceptors. Colored shaded area indicates the promoter (pink/yellow) or gene body region (purple) for each gene.

